# Orientation-selective adaptation improves perceptual grouping

**DOI:** 10.1101/519561

**Authors:** Noga Pinchuk-Yacobi, Dov Sagi

## Abstract

The role of visual pattern adaptation, and learning, in spatial integration was investigated. Observers reported whether a grid of identical tilted bars is perceived as rows or columns (perceptual grouping task). Performance was measured multiple times during a session to determine effects of repeated exposure to the stimuli. To test for possible effects of learning on the within-session dynamics, observers repeated the experiment on five days. We found that repeated performance produced rapid within-day improvements, which were largely transient, and were not retained when tested on subsequent days. In addition, exposure to stimuli with equal orientation contributed to the within-session improvement, whereas stimuli having an orientation differing by 45° from the original orientation diminished the improvement previously obtained in the same session. Practice with the task, over days, resulted in faster improvements. The transient nature of these exposure driven improvements and their susceptibility to interference by stimuli designed to reduce adaptation suggest that adaptation was their main cause. Finally, to investigate the effects of adaptation on internal noise and on spatial integration, we employed an external noise paradigm, showing that internal noise reduction resulted from adaptation. Internal noise was reduced only when spatial integration was effective, suggesting that adaptation improved perception of global stimulus properties. Overall, our results suggest that the grouping task benefits from an adaptation process that rapidly adjusts the visual system to the statistics of the visual stimuli. We suggest that this effect is achieved through spatial decorrelation of neural responses. With practice, those adjustments are made faster.

## Introduction

Grouping of spatially distinct elements into coherent objects, a fundamental function of the visual system, requires spatial integration, with its effectiveness depending on long-range correlations in the image. A previous report demonstrated the effectiveness of a model based on spatial-correlations to successfully account for performance in a perceptual grouping task (Ben-Av & Sagi, 1995). It was shown that for a task in which observers had to report the organization of the stimuli as either horizontal rows or vertical columns, grouping occurred in the direction with higher spatial-correlations. However, brain representations of the stimuli are contaminated with internal, neuronal, noise. Uncorrelated internal noise is averaged out by spatial integration, but spatially correlated internal noise may bias the grouping signal and distort perception. It was demonstrated that even weakly correlated noise substantially limits effective signal capacity, which in return can reduce psychophysical performance (Zohary, Shadlen, & Newsome, 1994).

Visual adaptation is considered here as a stimulus-driven process which continuously and quickly adjusts the neural responses to the statistics of the current visual environment. Contrary to perceptual learning, which leads to long-term improvements in performance, adaptation effects are typically short-termed. Previous studies tested the effects of visual pattern adaptation on visual sensitivity. Adaptation to intensity-modulated gratings typically shows reduced sensitivity to low-contrast gratings of similar orientation and spatial frequency (Blakemore & Campbell, 1969). Some evidence for improved contrast discrimination around the adapted contrast level was provided, but these effects were small and inconsistent (Barlow, Macleod, & Van Meeteren, 1976; Greenlee & Heitger, 1988; Määttänen & Koenderink, 1991; Ross, Speed, & Morgan, 1993). Similarly, only mixed and weak results were found for improved discrimination following face adaptation (Ng, Boynton, & Fine, 2008; Rhodes, Maloney, Turner, & Ewing, 2007; Yang, Shen, Chen, & Fang, 2011). Sensory adaptation has been proposed to improve coding efficiency by reducing redundancy in sensory signals, possibly by decorrelating neuronal responses (Barlow & Földiák, 1989). Experimental support for this idea were provided by studies showing that adaptation decorrelates neural responses in monkey V1 neurons (Gutnisky & Dragoi, 2008), and maintains decorrelation across the population of cat V1 neurons (Benucci, Saleem, & Carandini, 2013). Thus, spatial decorrelation is expected to improve performance on visual tasks involving spatial integration, such as perceptual grouping.

Here we examined the spatial aspects of visual adaptation, by testing the effects of adaptation on the spatial integration of visual information. For this purpose, we tested how the performance of a perceptual grouping task, which includes repeated exposure to the grouping stimulus, changes during a testing session. To test the possible effects of experience with the stimuli on the within-session dynamics (Yehezkel, Sagi, Sterkin, Belkin, & Polat, 2010), observers performed five daily sessions. We found that repeated performance of the grouping task produced a rapid within-day improvement, which was largely transient, and was not fully retained when tested on subsequent days. We took the transient nature of the gains to imply that they resulted from rapid adaptation to the visual stimuli. Practice with the task, over several days, resulted in faster adaptation, in accordance with previous findings (Yehezkel, Sagi, Sterkin, Belkin, & Polat, 2010). To further test our suggestion that exposure-based adaptation affects the performance in the grouping task, we explicitly tested how performance on the task changed following exposure to stimuli with the same orientation as in the grouping task or to stimuli with the orientation offset by 45°. Performance in the task improved following adaptation to the same stimuli, whereas adaptation to stimuli with orientation offset by 45° (de-adaptation) reduced performance. Finally, to uncover the effects of adaptation on internal noise and on noise reduction, we employed an external noise paradigm (Green & Swets, 1966; Lu & Dosher, 2008). The results showed reduced internal noise as a consequence of adaptation, with learning reducing the effect of external noise.

Overall, our results suggest that the grouping task benefits from an adaptation process that rapidly adjusts the visual system to the statistics of the stimulus orientations. With practice, this adjustment can be made faster.

## Methods

### Apparatus

The stimuli were presented on a 23.6” VIEWPixx/3D monitor (1920 × 1080, 10bit, 120Hz, with ‘scanning backlight mode’) viewed at a distance of 100 cm. The mean luminance of the display was 47.26 cd/m^2^ in an otherwise dark environment.

### Observers

Thirty-five observers with normal or corrected-to-normal vision participated in the experiments. All observers were naïve to the perceptual grouping task and gave their written informed consent. The work was carried out in accordance with the Code of Ethics of the World Medical Association (Declaration of Helsinki).

### Stimuli and task

#### Perceptual grouping task

The stimulus, of 40 ms duration, consisted of a grid of identical diagonal bars (19 ×19, tilted 45° counterclockwise from the vertical, 26.4 × 2.8 arcmin, length × width, and spaced 46.8 arcmin apart). The experimental variable was the difference between the horizontal spacing and the vertical spacing between the bars (dh-dv), thus controlling proximity-based grouping (Ben-Av & Sagi, 1995). The 13 differences tested were ±13.9, ±10.8, ±7.4, ±3.9, ±2.0, ±1.0 and 0 arcmin. A positive difference implies smaller spacing in the vertical direction, a columnar organization. Since the unequal spacing between the rows and the columns also affected the overall shape of the stimulus, introducing unwanted cues, a circular frame was added (radius= 366 arcmin), so that the global form of the stimulus was circular across all experiments (Fig. 1A). A 2AFC (2-Alternative-Forced-Choice) task was used, where observers reported the perceived organization of the display as one of two possible configurations: rows or columns. Response feedback was provided only during the familiarization session performed on the first day (to be consistent with previous studies using the same stimuli and task, e.g. Yehezkel et al., 2010). Each block of trials, of about 4 minutes duration, contained 12 or 13 dh-dv differences (depending on the inclusion of the dh-dv=0), intermixed (8 or 9 trials per difference). The measured psychometric curves (the percentage of vertical responses as a function of dh-dv) were fitted with a cumulative normal distribution (Fig. 2). The parameters obtained were used to estimate the point of subjective equality (PSE) where the rows and columns were reported with equal probability, and the slope of the psychometric function at that point (performance sensitivity). The discrimination threshold was defined as the standard deviation (σ) of the normal distribution fitted to the produced psychometric functions (lapse rate: up to 0.05, separately for lambda and gamma). Fitting was performed using Psignifit 4.0 software for MATLAB (Schütt, Harmeling, Macke, & Wichmann, 2016). In order to verify that the grid stimulus did not cause a specific bias in favor of one of the perceptual organizations (horizontal or vertical), we calculated for each observer the average PSE (across all blocks). The average PSE across observers was not significantly different from zero (0.28±0.4, mean±SEM, p=0.5, paired t-test), indicating that there was no consistent perceptual bias in favor of one of the perceptual organizations.

**Fig. 1.**
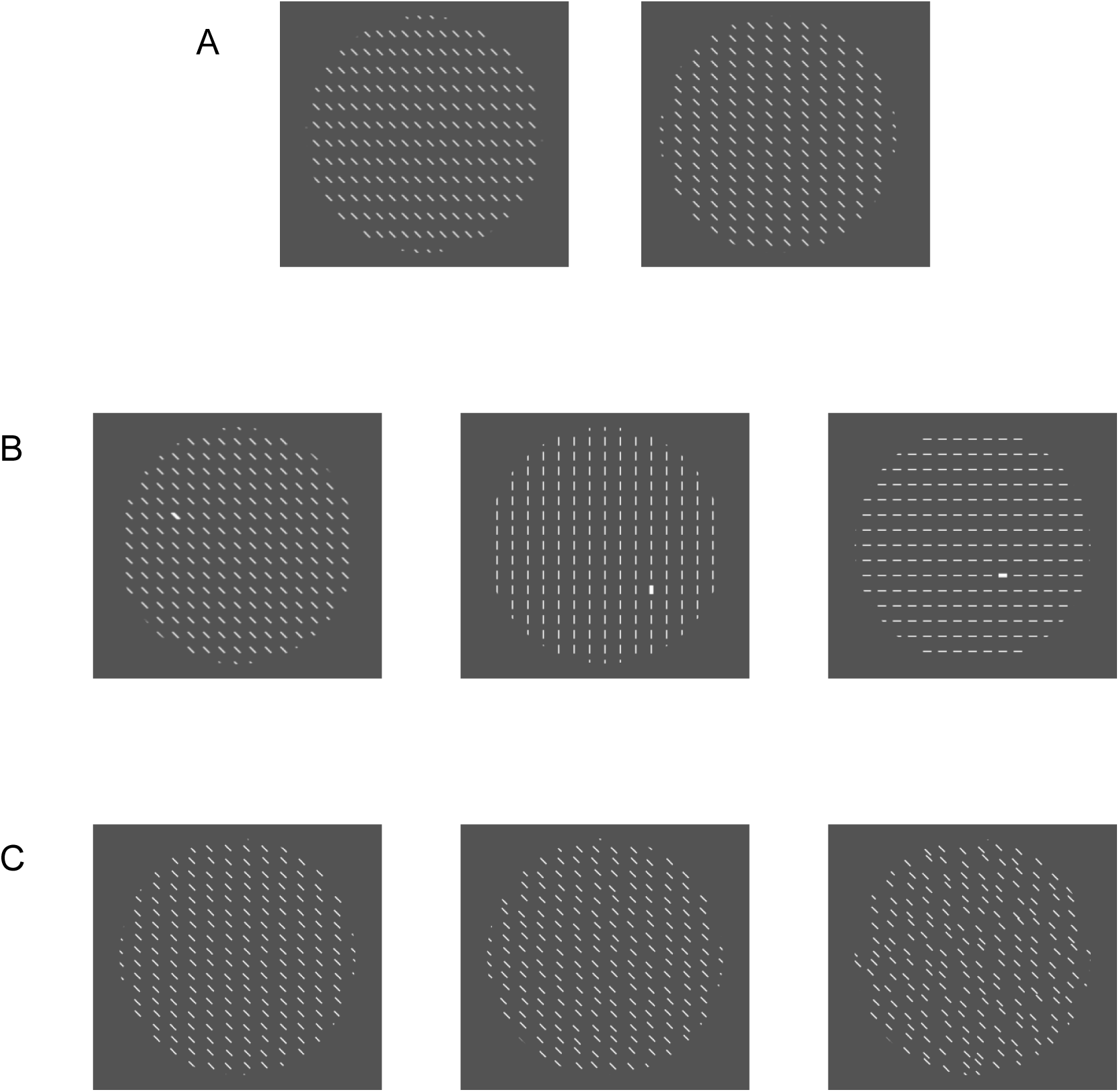
Stimuli and task. (A) Perceptual grouping task: The stimulus consisted of a grid of identical tilted bars (tilted 45° counterclockwise from the vertical, 40 ms duration) embedded within a circular window. The difference between the horizontal and vertical distances (dh-dv) between the bars changed from trial to trial. Observers were requested to report whether the stimulus is grouped into horizontal rows (left image) or vertical columns (right image). (B) Adaptation induction: The stimulus was the same grid of bars as in the grouping task, repeatedly presented for 40 ms. In order to ensure that observers were indeed looking at the stimuli, they performed an easy pop-out detection task, in which they reported whether a wider bar was presented on the left side or on the right side of the stimulus. There were three adaptation blocks types: 1.’Diagonal’ blocks that contained diagonal bars, the same as in the perceptual grouping task (left image). 2.’Vertical’ blocks that contained vertical bars (central image). 3.’Mixed’ blocks, in which the orientation of the bars changed randomly from trial to trial, and was either vertical or horizontal (central and right images). (C) Stimuli with external noise: low noise (left image), intermediate noise (central image), and high noise (right image).

**Fig. 2.**
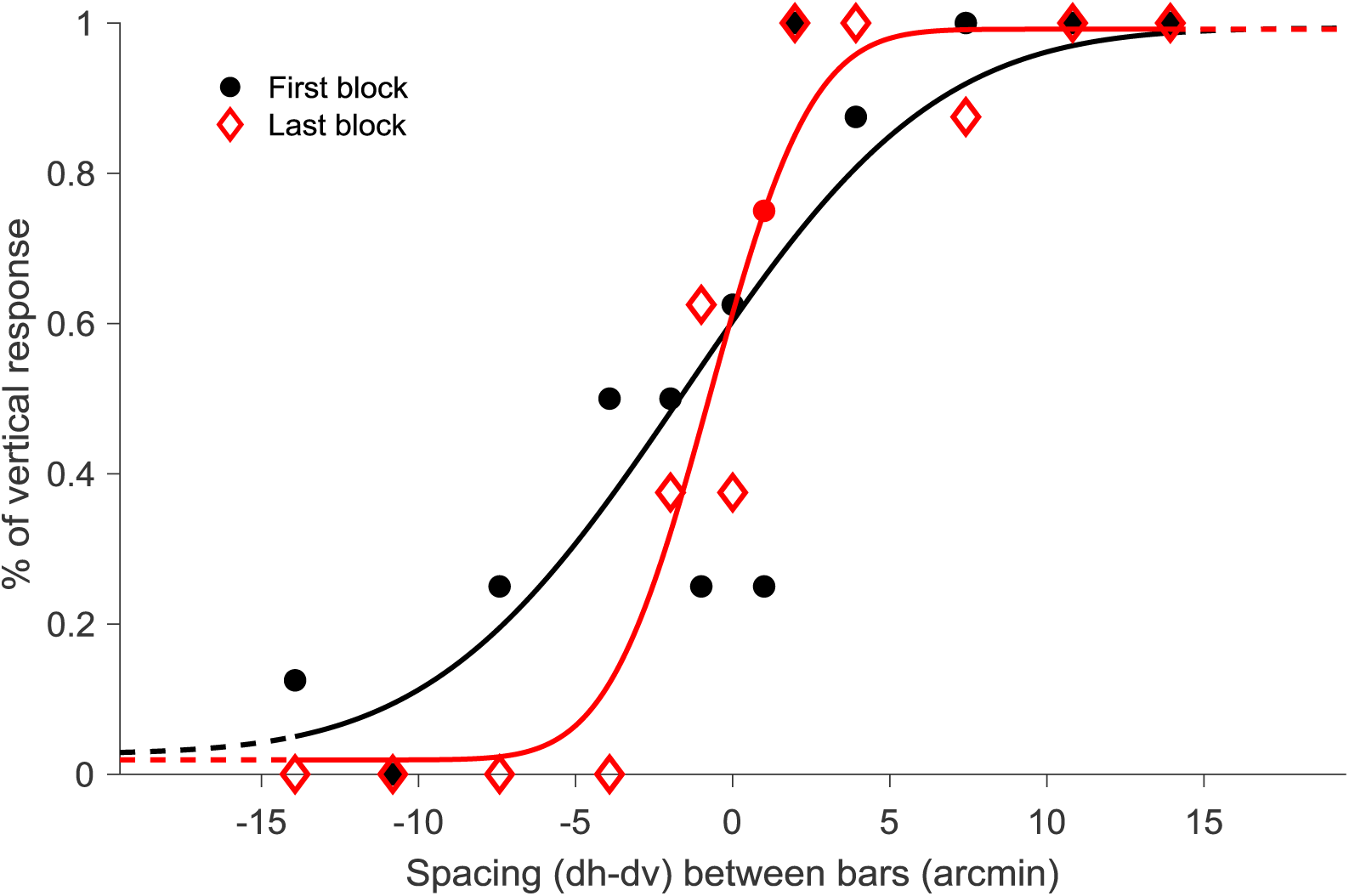
A pair of psychometric functions (percentage of vertical responses as a function of dh-dv), fitted with the cumulative normal distribution. The slope at the last block of the session is steeper, relative to the first block, showing improved sensitivity (N=8 trials/measurement).

#### Adaptation induction

Observers were presented with trials exposing the same grid of bars as in the grouping task (40ms duration), but without performing the task. To ensure that the observers were effectively exposed to the stimuli, they were asked to report the presence/absence of an easily detectable (>90% correct detection) wide bar, presented in the left or right stimulus quadrant with equal probability (Fig. 1B). Each adaptation block contained 5 dh-dv differences (±2.0, ±1.0 and 0 arcmin), intermixed (40 trials per difference), and lasted about 4 min. There were three types of adaptation blocks: 1. ‘Diagonal’ blocks in which the orientation of the bars in the adapting stimulus was the same as in the perceptual grouping task (tilted 45° counterclockwise from the vertical). 2. ‘Vertical’ blocks, in which the orientation of the adapting stimulus was vertical. 3. ‘Mixed’ blocks, in which vertical and horizontal adapting stimuli were randomly mixed during adaptation (Fig. 1C). Exposure to stimuli having the same orientation as the tested grouping stimuli (‘Diagonal’ stimuli) is expected to enhance adaptation to the grouping stimuli, whereas exposure to stimuli with 45° offset orientation (‘Vertical’ and ‘Mixed’ stimuli) is expected to reduce adaptation to the previous orientation (Greenlee & Magnussen, 1988; Harris, Gliksberg, & Sagi, 2012).

### Procedures

#### Experiment 1: Testing perceptual grouping

Observers (N=12) carried out the perceptual grouping task during five daily sessions (four blocks on day 1, five blocks on days 2-3, and six blocks on days 4-5). The time interval between subsequent sessions was 1-4 days. Overall, observers performed 2704 grouping trials (26 blocks × 13 dh-dv differences × 8 trials per difference).

#### Experiment 2: Adaptation and de-adaptation

Two experiments examined the dependence of the grouping performance on the adaptation/de-adaptation induction.

##### Experiment 2a

Nine of the observers that participated in the first experiment participated in this experiment, which included four daily sessions. In each session, observers were tested with three blocks of the perceptual grouping task, followed by three blocks of adaptation/de-adaptation, and then tested again with three blocks of the perceptual grouping task. Two of the sessions contained ‘Diagonal’ adaptation blocks (the same orientation as in the perceptual grouping task), whereas the other two sessions contained either ‘Vertical’ de-adaptation blocks (six observers) or ‘Mixed’ de-adaptation blocks (three observers). Overall, each observer performed 2496 grouping trials (24 blocks × 13 dh-dv differences × 8 trials per difference) and 2400 adaptation/de-adaptation trials.

##### Experiment 2b

Ten new observers performed the grouping task in testing blocks that were interleaved with adaptation/de-adaptation blocks. The experiment included two daily sessions, each containing five blocks of the perceptual grouping task and four adaptation/de-adaptation blocks (‘Diagonal’ or ‘Mixed’, respectively). The Grouping blocks were interleaved with the adaptation/de-adaptation blocks to create one of two block sequences: G→Ad→G→deAd→G→Ad→G→deAd→G or G→deAd→G→Ad→G→deAd→G→Ad→G, where G denotes a grouping block, Ad denotes a ‘Diagonal’ adaptation block, and deAd denotes a ‘Mixed’ de-adaptation block. Observers performed both sequences, on different days, starting with either the first sequence or the second sequence, counterbalanced between observers. Overall, each observer performed 1040 grouping trials (10 blocks × 13 dh-dv differences × 8 trials per difference) and 1600 adaptation/de-adaptation trials.

#### Experiment 3: External noise

Observers performed the perceptual grouping task with external noise added to the stimuli. External noise was added as a random uniformly distributed displacement (in x and y directions) to the position of each element in the display. There were two groups, either the ‘lower noise’ group (N=7) or the ‘higher noise’ group (N=6), differing in their levels of external noise included. There were five daily sessions, each with two blocks of trials. Each block contained equal numbers of randomly intermixed trials that included stimuli either without external noise or with one of two noise levels. In the ‘lower noise’ group, the external noise was either low or intermediate (uniformly distributed within a range of ±2.8 or ±5.6 arcmin). In the ‘higher noise’ group, the external noise was either intermediate or high (uniformly distributed within a range of ±5.6 or ±11.2 arcmin). Overall, the observers performed 3240 trials (10 blocks × 12 dh-dv differences × 9 trials per difference × 3 noise levels).

##### Modeling the external noise

Testing human sensitivity to external noise allows one to characterize the limiting properties of task performance, arising from internal and external noise (Lu & Dosher, 1999; Nagaraja, 1964). We fit the measured thresholds to a standard model assuming the additivity of external noise and equivalent internal noise, the latter representing internal variations expressed in terms of stimulus variations, limiting performance in the absence of external noise. Accordingly, we defined threshold as the displacement (dh-dv) value that is equal to the total noise in the stimulus, that is, the sum of variations generated by the external noise (sum of the noise variances in the X and Y direction) and the equivalent internal noise. This threshold was estimated by the standard deviation (σ) of the normal distribution fitted to the produced psychometric functions (see above, stimuli and task). We further assumed that the visual system integrates the spatially distributed external noise, with an integration coefficient that depends on the adaptation level and experience with the task. This model is summarized by the following equation:

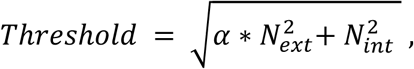

where α denotes the (inverse) efficiency of external-noise integration, estimating the capacity of the grouping process to reduce the impact of external noise by spatial integration. N_int_ and N_ext_ correspond to the SD of the equivalent-internal and external noises, respectively, and both are expressed in stimulus units. Note that we do not assume that the efficiency parameter controls the effective internal noise, an assumption that requires the internal noise to be spatially uncorrelated. We tested the dependence of the two free parameters, α and N_int_, on experience within and between sessions, by fitting the above equation to the threshold vs external noise data at each session start and end, for each observer separately. In each such fit, there were 3 external noise levels (zero, low and intermediate levels in the ‘lower noise’ group; zero, intermediate and high in the ‘higher noise’ group) and 3 threshold measurements. A least square fitting method was used. Note that α and N_int_ may display similar dependencies on time, since N_int_ may improve with spatial integration, depending on spatial correlations within the visual system.

## Results

### Experiment 1: Repeated performance on the perceptual grouping task

The observer’s performance on the perceptual grouping task was measured by the percentage of ‘vertical’ reports as a function of dh-dv (psychometric function, Fig. 2). The discrimination threshold was defined as the standard deviation of the normal distribution fitted to the produced psychometric functions (see Methods). Figure 3 displays the average discrimination threshold (across observers) for each testing block and daily session. Results show significant within-session improvements, indicated by a significant decrease in the threshold from the first training block to the last training block (2.5±0.6, p<0.01; 3.1±0.9, p<0.01; 3.0±0.8, p<0.01; 1.1±0.6, p=0.09; 2.0±0.6, p<0.01; mean±SEM, paired t-test, improvements within 1^st^, 2^nd^, 3^rd^, 4^th^, and 5^th^ sessions, respectively). In addition, the first threshold measured on each day was significantly higher than the threshold of the last training block of the previous day (2.5±0.6, p<0.01; 2.3±0.6, p<0.01; 0.9±0.4, p<0.05; 1.7±0.3, p<0.01; mean±SEM, paired t-test, threshold increase between sessions 1-2, 2-3, 3-4, and 4-5, respectively), showing that the within-day performance gains were not fully retained in subsequent daily sessions. The transient nature of these gains suggests the involvement of visual adaptation as their main cause.

**Fig. 3.**
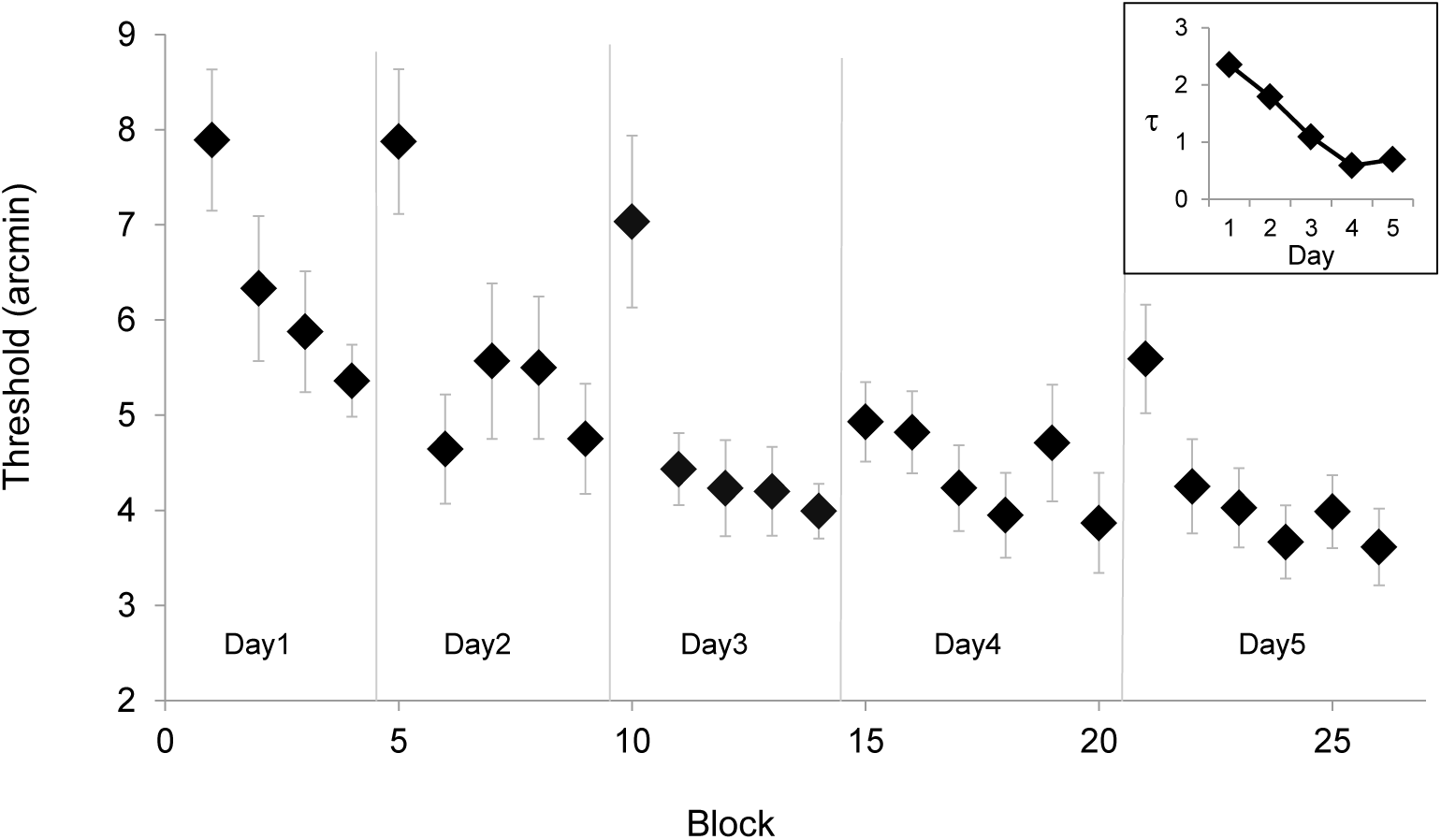
Grouping thresholds in the perceptual grouping task, showing within-day improvements that are not fully retained in subsequent daily sessions. Thresholds are averages across observers (N=12) with error bars corresponding to ±1SEM. The inset plot displays the time constant (τ sessions) of an exponential decay model fitted to the daily thresholds, showing that the within-session learning rate improves from day to day (see text for details).

Next, we investigated whether the dynamics of performance changed between days. The results show that there were no significant changes between the threshold reductions within the first day and the threshold reductions within subsequent days (first vs second day: p=0.5, first vs third day: p=0.6, first vs fourth day, p=0.2, first vs fifth day: p=0.5, paired t-test). However, as shown in Fig. 3, practice with the task resulted in faster within-session threshold reductions. On day 1 the threshold was gradually reduced. On days 2 and 3, the threshold was sharply reduced already in the second block. Whereas the threshold of the first block in days 2 and 3 was similar to the threshold of the first block in the first day (1st day: 7.9±0.7, 2nd day: 7.9±0.8, p=1.0, 3rd day: 7.0±0.9, p=0.5, mean±SEM, paired t-test), the threshold of the second block in these days was significantly reduced as compared to the threshold of the second block in the first day (1st day: 6.3±0.8, 2nd day: 4.6±0.6, p<0.01, 3rd day: 4.4±0.4, p<0.01, mean±SEM, paired t-test). With more practice, the threshold reduction on days 4 and 5 started even faster, already in the threshold of the first block, which was significantly reduced compared to the threshold of the first block of the first day (4th day: 4.9±0.4, p<0.01, 5th day: 5.6±0.6, p<0.05, mean±SEM, paired t-test). To quantify the speedup of the within-session threshold decrease, we fitted the daily thresholds (averaged across observers) to an exponential decay model, assuming that the initial and final thresholds are fixed across days. The model was successful in accounting for 86% (R^2^) of the variance in the data (Fig. 3: 26 datum points, 7 free parameters). We found that the time constant gradually decreased during the 5 testing days, showing ∼4 times faster within session threshold reduction at day 5 as compared with day 1 (see inset, Fig. 3). These faster within-session improvements with practice could possibly result from faster re-adaptation (Yehezkel et al., 2010).

### Experiment 2: Adaptation/de-adaptation

Here we set out to test the hypothesis that the transient within-day improvement obtained in the grouping task was due to task-independent, exposure based, sensory adaptation.

#### Experiment 2a

Nine observers from Experiment 1 were exposed to trials presenting the grouping stimuli, but they performed a detection task instead of the grouping task (see Methods). Figure 4 displays the average grouping performance in daily sessions that contained ‘Diagonal’ adaptation blocks, and in daily sessions that contained ‘Vertical’ or ‘Mixed’ de-adaptation blocks. The results show that following exposure to stimuli having the same orientation as the grouping stimuli (‘Diagonal’), the within-session improved grouping performance was preserved, and somewhat improved, as expected from sensory adaptation. However, with the ‘Vertical’ and ‘Mixed’ exposure blocks, the grouping thresholds significantly increased, in agreement with the de-adaptation hypothesis. In order to investigate whether these trends were statistically significant, we quantified for each observer the pre-exposure performance and the post-exposure performance. The pre-exposure performance was calculated as the average threshold of the two perceptual grouping blocks before exposure, and the post-adaptation performance was calculated as the average threshold of the two blocks after exposure. The differences between the pre-exposure and the post-exposure performances were statistically significant for the different adaptation/de-adaptation types (‘Diagonal’ adaptation: −0.8±0.2, ‘Vertical’ & ‘Mixed’ de-adaptation: 0.7±0.2, pairwise t-test between conditions showing p<0.01, Fig. 4). The threshold was significantly reduced following exposure to ‘Diagonal’ blocks (pre-adaptation: 4.3±0.4, post-adaptation: 3.5±0.4, p<0.05, paired t-test, Fig. 4), and significantly increased following exposure to ‘Vertical’ & ‘Mixed’ blocks (pre-adaptation: 3.98±0.3, post-adaptation: 4.64±0.4, p<0.05, paired t-test, Fig. 4).

**Fig. 4.**
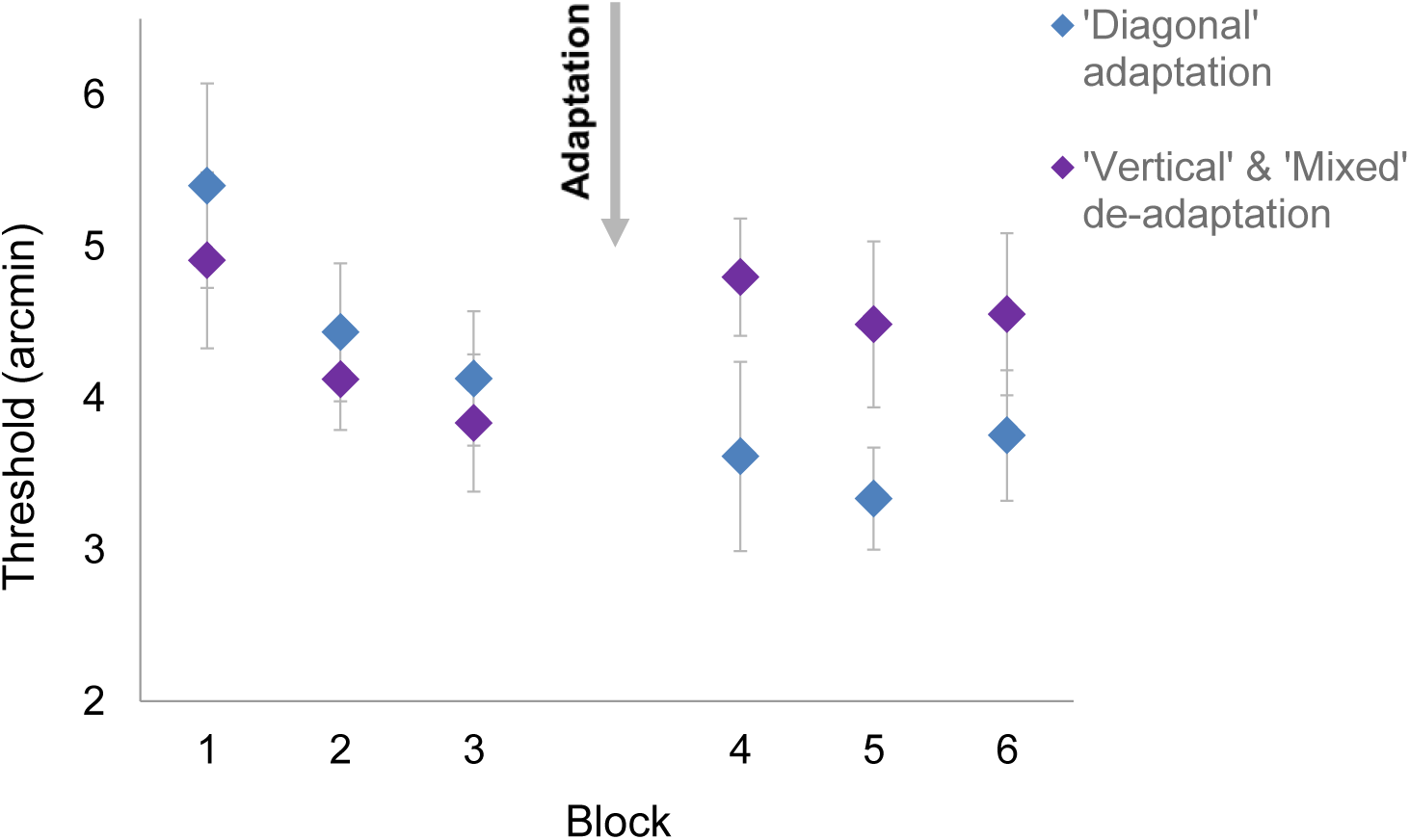
Grouping threshold obtained in daily sessions that contained adaptation to stimuli with ‘Diagonal’ orientation (the same orientation as in the grouping task, blue rhombuses), and daily sessions that contained de-adaptation to ‘Vertical’ & ‘Mixed’ stimuli (purple rhombuses). Thresholds are averages across observers (N=9) with error bars corresponding to ±1SEM.

#### Experiment 2b

Ten new observers performed grouping blocks that were interleaved with exposure blocks. Figure 5 displays the average grouping performance (across observers) for the two interleaved block sequences. The average threshold (across blocks) following a ‘Diagonal’ exposure (adaptation: 4.8±0.5, SEM) was significantly lower (p<0.01, paired t-test) than the average threshold following a ‘Mixed’ exposure (de-adaptation: 6.1±0.8, SEM). Here we used only the ‘Mixed’ blocks to avoid possible appearance biases that resulted from continuous adaptation to vertical lines as in the ‘Vertical’ blocks.

**Fig. 5.**
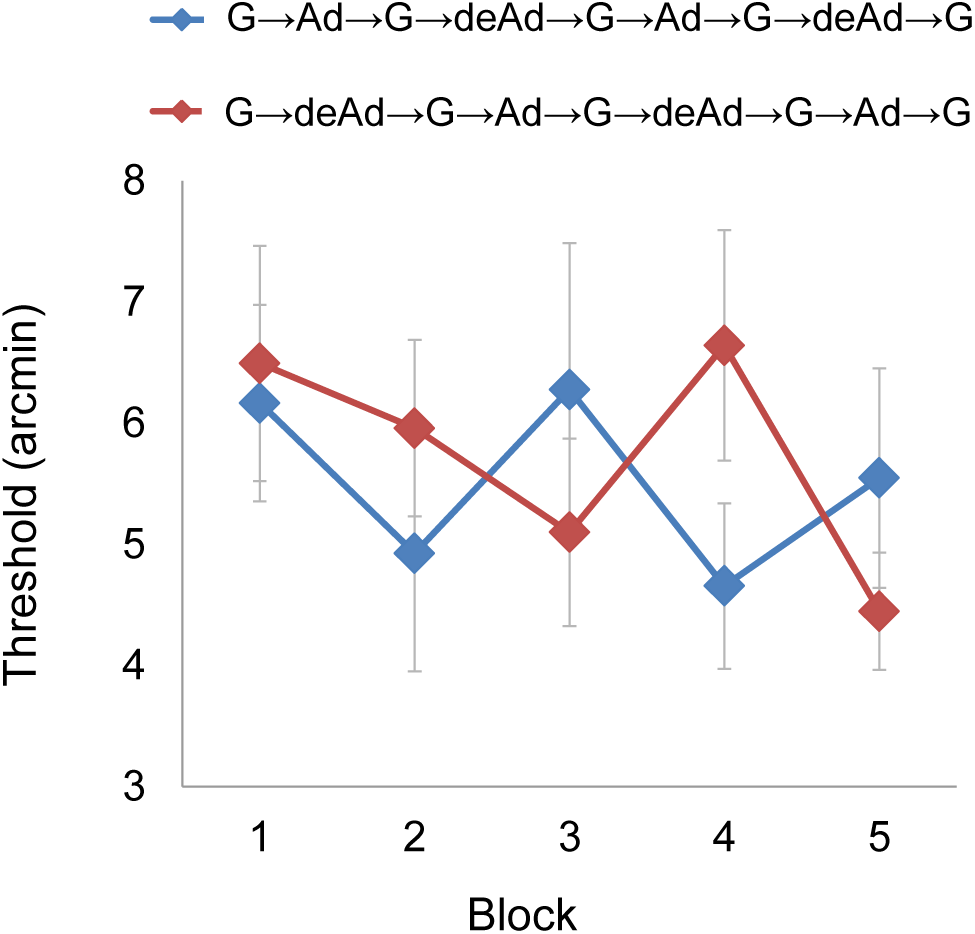
Grouping thresholds obtained in sessions that contained the G→Ad→G→de-Ad→G→Ad→G→de-Ad→G block sequence (blue rhombuses) and sessions that contained the G→deAd→G→Ad→G→de-Ad→G→Ad→G block sequence (red rhombuses). G denotes a grouping block, Ad denotes a ‘Diagonal’ adaptation block and deAd denotes a ‘Mixed’ de-adaptation block. Note that there was no threshold elevation following the first de-adaptation block (second red rhombus), probably because one grouping block is not of sufficient length to cause threshold reduction due to adaptation (supported by results from experiment 1, Fig. 3), therefore there was nothing for the de-adaptation block to eliminate. Thresholds are averages across observers (N=10) with error bars corresponding to ±1SEM.

The results of the two adaptation/de-adaptation experiments support the hypothesis that the transient improvements at the beginning of each session result from a task-independent adaptation process that rapidly adjusts the visual system to the stimulus orientation. The improved performance following exposure to stimuli with the same orientation is suggested to arise from adaptation, since perceptual learning is thought to be task specific (Ahissar & Hochstein, 1993; Fahle & Morgan, 1996). Most importantly, exposure to stimuli with 45° offset orientation, as in the ‘Vertical’ and ‘Mixed’ blocks, resulted in a significant deterioration of performance, rather than improvement. This result is in accordance with the adaptation literature showing de-adaptation for stimuli offset by 45° (Greenlee & Magnussen, 1988; Harris et al., 2012; Pinchuk-Yacobi, Harris, & Sagi, 2016)(Greenlee & Magnussen, 1988; Harris et al., 2012).

### Experiment 3: Effects of external noise

Here we tested the performance in the task while adding different levels of external noise to the stimuli. Figure 6A and 6B display the average discrimination threshold (across observers) for each testing block and noise level, for the ‘lower noise’ group and the ‘higher noise’ group, respectively.

**Fig. 6.**
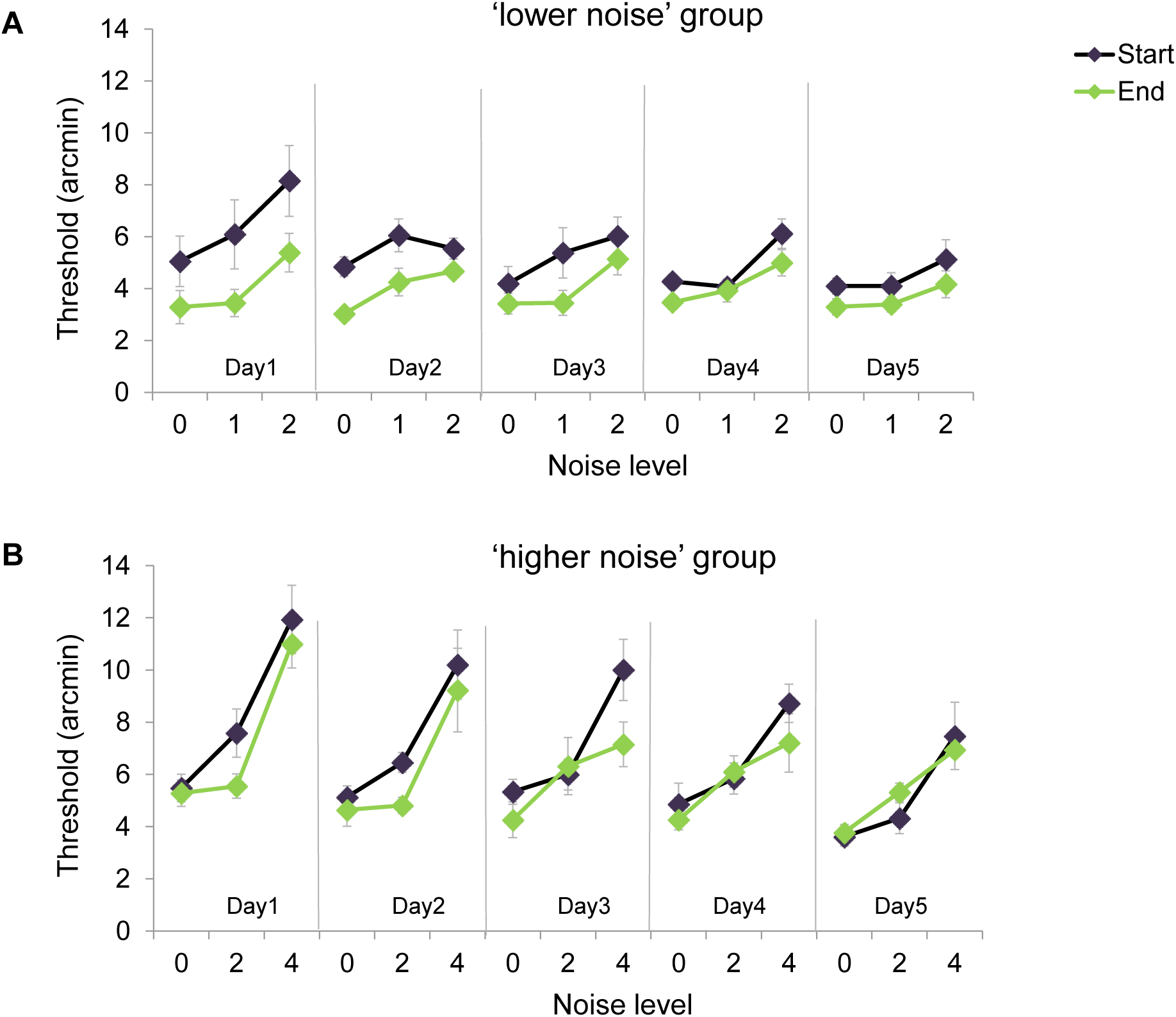
Performance in the perceptual grouping task with external noise. The experiment included five daily sessions, with two blocks per day (Start and End). Each block contained equal numbers of randomly intermixed trials with stimuli either without external noise (noise level =0) or with one of two noise levels. Overall, there were three external noise levels: low (‘1’, jitter of 2.8 arcmin), intermediate (‘2’, jitter of 5.6 arcmin), and high (‘4’, jitter of 11.2 arcmin). (A) Performance of the ‘lower noise’ group (N=7), low and intermediate noise levels. (B) Performance of the ‘higher noise’ group (N=6), intermediate and high noise levels. Thresholds are averages across observers with error bars corresponding to ±1SEM.

The effects of experimental manipulations on threshold were evaluated using a repeated measures ANOVA with the within-observers factors including noise (2 levels: no noise and mid noise levels), day (5 days), and time (start and end of session) performed separately for the two groups. We used only the data obtained from trials with no noise and mid noise levels, since only those noise levels were used in both groups (see Appendix for statistical analysis performed on the whole data). For both groups, threshold increased significantly when external noise was added to the stimuli (F(1,6) = 130.3, p< 0.001, F(1,5) = 10.3, p<0.05, the ‘lower noise’ group and the ‘higher noise’ group, respectively). Improvement within session, as indicated by the threshold reduction from the start of the session to the end of the session, was only significant for the ‘lower noise’ group (F(1,6) = 93.6, p< 0.01, F(1,5) = 3.7, p =0.1, the ‘lower noise’ group and the ‘higher noise’ group, respectively).

In order to test the effect of learning in the task, we ran an ANOVA with only the thresholds of the first day (day1) and the last day (day 5), separately for the thresholds at the start and at the end of the session. Learning between the first day and the last day was only significant for the ‘higher noise’ group, and only at the start of the session (start: F(1,6) = 3.7, p =0.1, F(1,5) = 10.3, p< 0.05; end: F(1,6) = 0.06, p =0.8, F(1,5) = 5.9, p =0.06, the ‘lower noise’ group and the ‘higher noise’ group, respectively).

Next, we investigated whether the within-session gains, obtained in a training session, were retained in the following session, or whether performance deteriorated from the end of the session to the beginning of the next session. For that purpose, we ran another repeated measures ANOVA with the within-subject factors noise (no noise and mid noise levels), day (day 2-day 5), and time (end of the previous session or the start of the next session). Performance significantly deteriorated from the end of the previous session to the beginning of the next session only for the ‘lower noise’ group (F(1,6) = 31.1, p =0.001, F(1,5) = 0.03, p =0.9, the ‘lower noise’ group and the ‘higher noise’ group, respectively). This was clearly seen on the 2^nd^ day (Fig. 6A), where noise levels 0 and 1 at the start of day 2 are back to the starting level of the first day.

To compare the performances of the groups, we added to the repeated measures ANOVAs the between-subject factor of group type (‘lower noise’ group or ‘higher noise’ group). The only significant differences between the groups were the improvements within-session (F(1,11) = 16.3, p< 0.01) and the deterioration between sessions (F(1,11) = 11.1, p< 0.01), which were significantly lower for the ‘higher noise’ group. These results suggest that mixing trials with high external noise, as with the ‘higher noise’ group, reduces the adaptation effect. The learning effects (start: F(1,11) = 1.0, p = 0.4; end: F(1,11) = 0.7, p = 0.4) and the noise effect (F(1,11) = 3.1, p = 0.1) were not significantly different between the groups. No significant interactions were found in any ANOVA analysis.

Table 1 summarizes the statistical results for the analysis performed on data only from trials with no noise and mid noise levels.

**Table 1.**
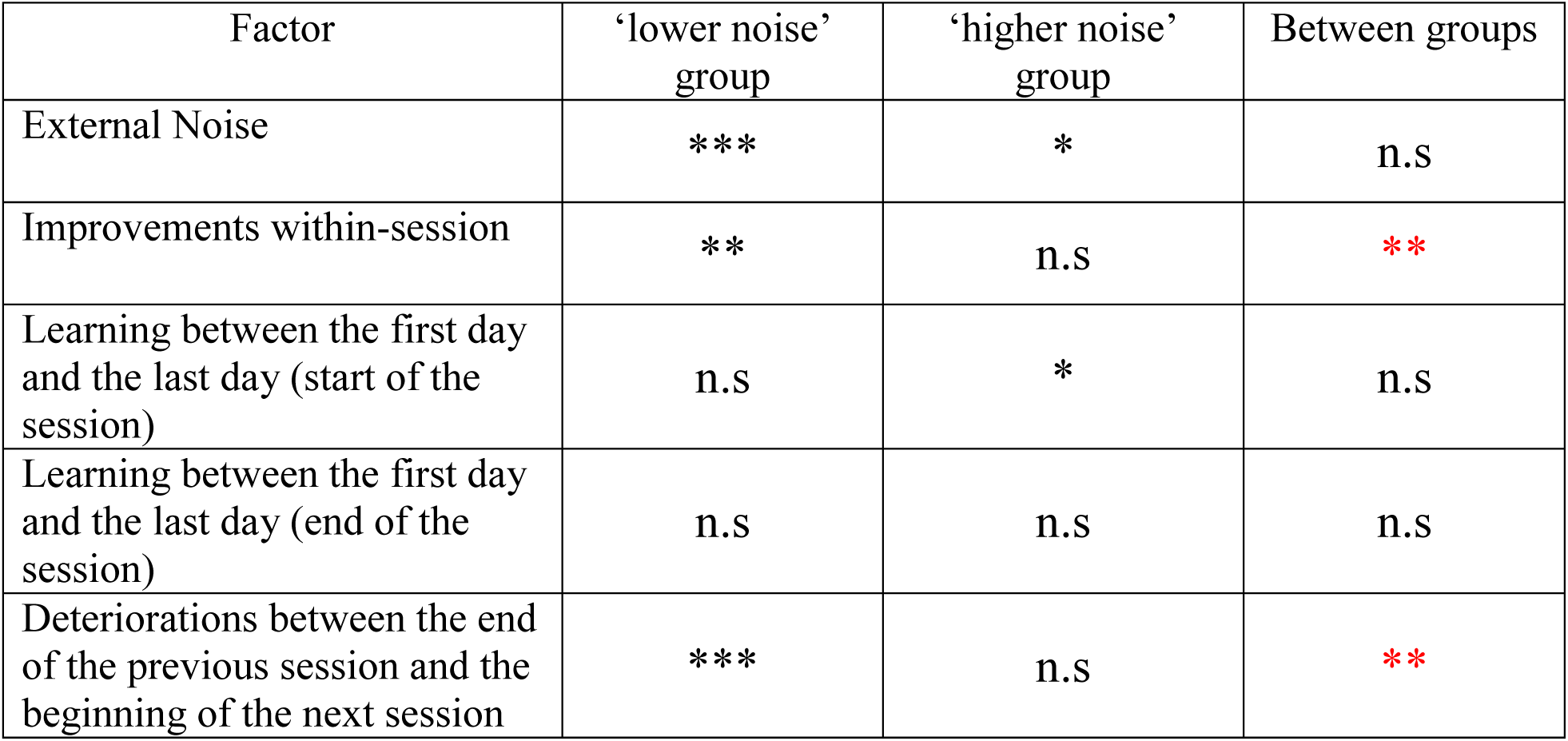
Statistical results for the external noise experiment. Analysis was performed only on data from trials with no noise and mid noise levels. *P < 0.05, **P < 0.01, ***P < 0.001.

In order to better explore the factors that could contribute to the changes between the groups’ thresholds, we fitted the thresholds (using data from both ‘lower’ and ‘higher’ noise groups) to the external-noise model that includes the parameters of an integration coefficient (α) as well as internal and external noise levels (see Methods). Fitting the integration coefficient resulted in inconsistent and minor changes (either a slight increase or a slight decrease) between subsequent sessions, that is, between a session end and the start of the following session. Therefore, we fixed the value of the integration coefficient between subsequent sessions (value at the start of the session equals to the value at the end of the previous sessions) to reduce the degrees of freedom. Results for the fitted parameters of the integration coefficient and the internal noise, averaged across observers, are displayed in Figs. 7A and 7B, respectively. The two parameters display different dependencies in the two tested groups. When only low and intermediate levels of external noise were added to the stimuli, as in the ‘lower noise’ group, observers quickly learned to integrate the bars, as indicated by a significant reduction of the integration coefficient, apparent already on the first training day (1.9±0.6, p<0.05, paired t-test). This fast improvement in integration, already on the first day, seemed to enable a significant within-session reduction in the internal noise (F(1,6) = 14.6, p< 0.01, two-way ANOVA with within-observers factors of day and time in the session), which was apparent already on the first day (2.0±0.6, p<0.05, paired t-test). The dependence of internal noise on spatial integration is supported by a significant correlation between the values of the two parameters (r=0.39, p<0.001, n=70). In contrast, the ‘higher noise’ group displays only a gradual reduction of the integration coefficient across days, and no significant reduction on the first day (0.3±0.2, p=0.2, paired t-test). Accordingly, there was no significant within-session reduction in internal noise (F(1,5) = 1.5, p = 0.3, two-way ANOVA with within-observers factors of day and time in the session), and there was no correlation between the values of the internal noise and the values of the integration coefficient (r=0.0, p=1.0, n=60). Comparing the two groups shows significantly lower internal noise at the end of the sessions in the ‘lower noise’ group compared with the ‘higher noise’ group (1.1±0.2, p<0.01, two-sample t-test). Given that the internal noise was significantly reduced only when spatial integration was present (as in the ‘lower noise’ group), suggests that adaptation does not improve performance by uniform reduction of the local internal noise, but by global decorrelation of internal noise across the whole stimulus.

**Fig. 7.**
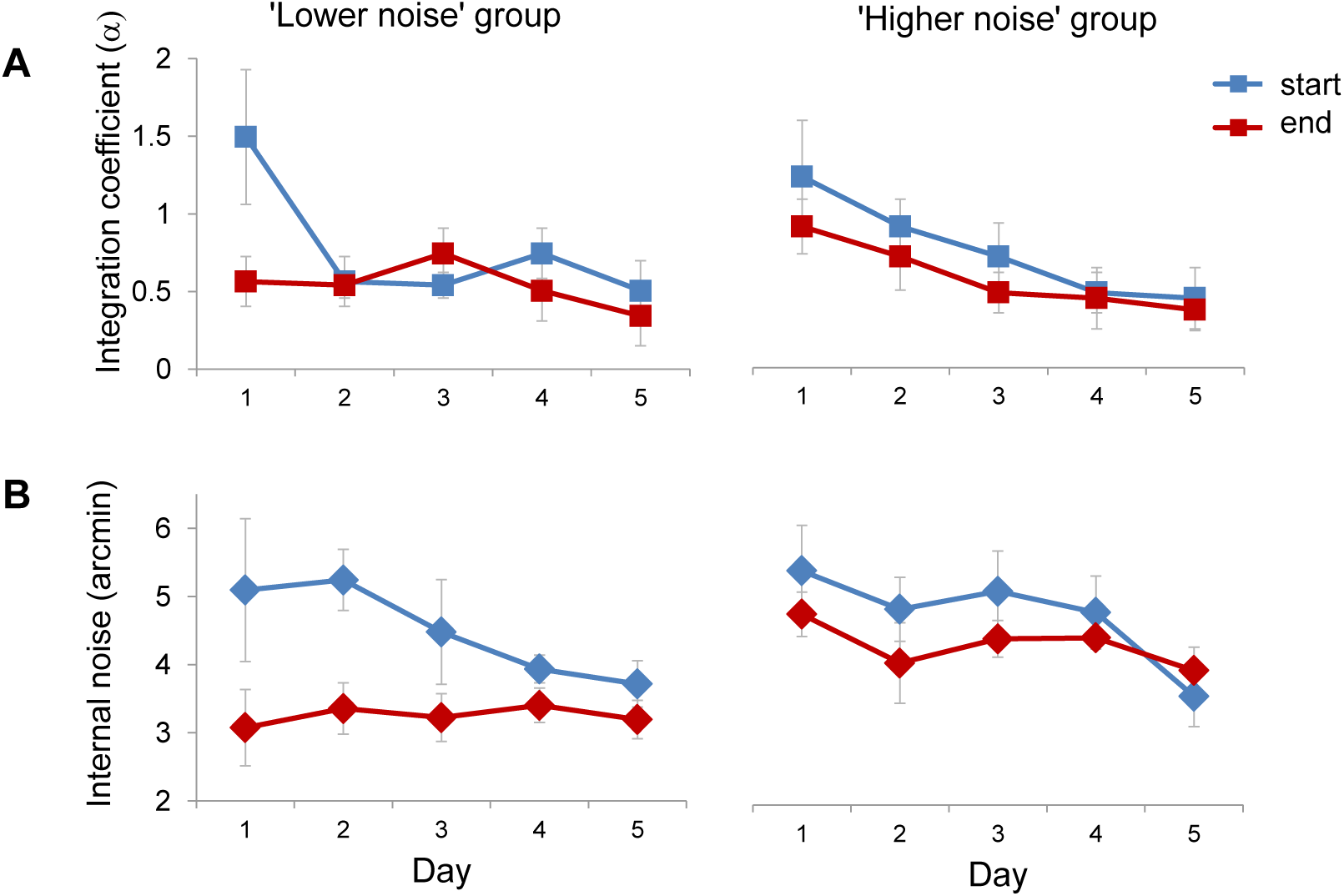
Fitted values of (A) the integration coefficient (α, lower value indicates improved spatial Integration), and (B) the internal noise, shown for each training day and time within sessions (start or end) for the ‘lower noise’ group (left, N=7) and for the ‘higher noise’ group (right, N=6). Results show that spatial integration is essential for obtaining lower internal noise. Error bars represent ±1SEM.

## Discussion

Our results show that repeated performance on the perceptual grouping task produced within-session improvements, especially at the beginning of each daily session (Fig. 3). These improvements were largely exposure (Fig. 4-5), not task dependent, and were not fully retained in subsequent daily sessions (Fig. 3). In addition, exposure to stimuli with an orientation differing by 45° from the orientation used in the grouping task diminished the improvement previously obtained in the same session (Figs. 4-5). The transient nature of these exposure-driven improvements, and their susceptibility to interference by de-adapting stimuli suggest that they result from an adaptation process that rapidly adjusts the visual system to the orientation statistics of the stimuli.

One of the main controversies regarding visual pattern adaptation concerns its potential benefit. At the single neuron level, adaptation was suggested to improve information transfer by adjusting the limited dynamic range of the neuron to the current input statistics (Wainwright, 1999). At the population level, adaptation was suggested to remove redundancies across neurons by de-correlating their neural responses (Barlow & Földiák, 1989). However, previous studies that tested how adaptation affects contrast discrimination found only weak evidence for improved discriminability for stimuli similar to the adapter, predicted by dynamic-range adjustment (Barlow et al., 1976; Greenlee & Heitger, 1988; Määttänen & Koenderink, 1991; Ross et al., 1993). Here we report large improvements in grouping discrimination following adaptation. The difference between our results and the results from previous perceptual adaptation experiments might arise from the different task we used. Perceptual grouping was suggested to rely highly on spatial correlations and spatial integration (Ben-Av, Sagi, & Braun, 1992), and therefore it should be most affected by adaptation-induced changes in spatial correlation. Contrast discrimination, on the other hand, shows no improvements from spatial integration when the base and increment contrasts are over an equal spatial extent, possibly due to balanced excitation and inhibition interactions (Bonneh & Sagi, 1999). Ben-Av & Sagi (1995) suggest that perceptual grouping involves a process that compares horizontal and vertical intensity correlations, and that grouping occurs in the direction with the higher degree of correlation. Given the high reliance of grouping on spatial correlations and integration, we argue that adaptation improves performance in the task by spatial decorrelation. Consequently, we propose two possible mechanisms for such improvements. One, removal of the average spatial correlation between all signals, assisting the comparison process in distinguishing between small differences in correlation values. Two, removal of correlations between the internal noises of neighboring neurons, increasing the efficiency of spatial integration and improving signal-to-noise ratio. An alternative explanation might have been that adaptation improves performance on the task by local means, such as reduction of internal noise at all locations, or local improvements in spatial or orientation resolutions. However, our results in the external noise experiment indicate that spatial integration is essential for obtaining the benefits of adaptation. Challenging spatial integration, by randomly mixing trials with high external noise as in the ‘higher noise’ group, resulted in reduced spatial integration and a significantly higher internal noise at the end of the sessions (Fig. 7). In accordance, the adaptation effects of within-session improvements and between-session deteriorations were significantly reduced in the ‘higher noise’ group (Fig. 6).

Comparing the performance in the perceptual grouping task in our experiments to the performance in other perceptual tasks shows compatible within-session improvements (Aberg, Tartaglia, & Herzog, 2009; Fahle, Edelman, & Poggio, 1995; Hussain, Sekuler, & Bennett, 2008; Karni & Sagi, 1993). However, contrary to other perceptual tasks in which the initial performance in each session is usually equivalent to (or even better than) the performance obtained at the end of the previous session (Harris & Sagi, 2015; Karni & Sagi, 1993), performance in the grouping task was largely reduced in the first block of subsequent sessions. This reduced initial performance indicates that the within-session gains obtained in the preceding training session were not retained in the next session (Fig. 3, the first three sessions) or were only partially retained (the fourth and fifth sessions). Failure to retain the within-session gains was also apparent in the repeated re-emergence of significant within-session improvements in all daily sessions. This behavior is inconsistent with typical within-session improvements obtained in other perceptual learning tasks, which mostly appear during the first training session and tend to diminish in subsequent sessions (Harris et al., 2012; Karni & Sagi, 1993). In addition to the diminished performance at the beginning of each session, which can be related to passive decay due to passage of time between sessions, performance within-session was also reduced after explicit exposure to de-adapting stimuli. Performance was diminished only following adaptation to stimuli that were offset by 45° from the orientation of the original grouping stimuli (‘Vertical’ or ‘Mixed’ orientation), and was enhanced following adaptation to stimuli with the same orientation as in the grouping task (‘Diagonal’ orientation). Thus, the deterioration effect was specific to the orientation of the stimuli. These results can be related to studies that tested the effects of adaptation on orientation mixtures. Greenlee & Magnussen (1988) found reduced contrast adaptation when vertical and diagonal gratings were interleaved during the adaptation period. Harris & Sagi (2015) found, using the texture discrimination task, slower within-session learning when using a fixed texture orientation, compared with conditions where two orientations differing by 45° were interleaved, attributing the effect to differences in orientation-specific adaptation.

Previous studies used external noise to explore the factors that contribute to the learning of a perceptual task (Lu & Dosher, 2008). Most relevant to our study is a study by Li, Levi, & Klein (2004) that used position noise to explore the neural mechanisms underlying learning of a visual position discrimination task. Li et al (2004) found that with learning observers improved performance by using more efficient and broader stimulus samples in their position judgments. These results are consistent with our findings of between-session learning of the integration coefficient. The within-session dynamics cannot be compared since performance results were only reported as within-session means. Furthermore, Li et al (2004) assumed internal noise to be spatially uncorrelated, thus predicting reduced effective internal noise with increasing sampling efficiency, while we suggest these correlations to be adaptation dependent. Interestedly, Li et al (2004) noted that, unlike with high external noise, in the low external noise condition, which was dominated by the internal noise, the measured human thresholds were much lower than the thresholds predicted by an ideal observer. This result can be explained by assuming that the internal noise contained correlations that limited the benefit obtained from more efficient and broader stimulus sampling.

The dynamics of the within- and the between-session gains in our experiment can also be related to studies of motor adaptation learning (Krakauer, 2009; Krakauer, Ghez, & Ghilardi, 2005). The return to the baseline performance level at the beginning of the second and third sessions resembles the ‘forgetting’ of motor improvements, when subjects are re-exposed to the same motor adaptation task later in time. In addition, the faster within-session threshold reduction with more daily sessions can be explained by a faster rate of re-adaptation, similar to faster relearning of a motor adaptation task following repetitive practice over time. Such faster re-adaptation was also shown recently following repeated adaptation to a prism distortion (Habtegiorgis, Rifai, Lappe, & Wahl, 2018; Yehezkel et al., 2010). These results of long-term effects of adaptation, which are found in several modalities, such as motor and vision, are consistent with theories suggesting that adaptation involve temporary plasticity of inhibitory synapses (Dealy & Tolhurst, 1974; Wilson, 1975). Perhaps when a substantial amount of adaptation is provided in a specific context (such as in the context of a perceptual task), the temporary plasticity can be made long-term.

## Acknowledgments

This work was supported by the Basic Research Foundation administered by the Israel Academy of Sciences and Humanities.

## Appendix

Here we performed the same ANOVA analyzes as described in the results section of Experiment 3, but on the whole data (including all noise levels). For both groups, thresholds increased significantly when external noise was added to the stimuli (F(2,12) = 40.3, p< 0.001, F(2,10) = 23.1, p< 0.001, the ‘lower noise’ group and the ‘higher noise’ group, respectively), and thresholds decreased significantly from the start of the session to the end of the session, demonstrating within-session improvements (F(1,6) = 72.8, p< 0.001, F(1,5) = 7.3, p< 0.05, the ‘lower noise’ group and the ‘higher noise’ group, respectively). Learning between the first day and the last day was only significant for the ‘higher noise’ group (start: F(1,6) = 3.2, p = 0.12, F(1,5) = 18.2, p< 0.01; end: F(1,6) = 0.007, p=0.9, F(1,5) = 22.6, p< 0.01, the ‘lower noise’ group, and the ‘higher noise’ group, respectively). Deterioration in performance from the end of the previous sessions to the beginning of the next session was only significant for the ‘lower noise’ group (F(1,6) = 45.7, p< 0.001, F(1,5) = 0.4, p = 0.6, the ‘lower noise’ group and the ‘higher noise’ group, respectively).

Table 2 summarizes the statistical results for the external noise experiment (analysis on the whole data).

**Table 2.** Statistical results for the external noise experiment. Analysis was performed on the whole data. *P < 0.05, **P < 0.01, ***P < 0.001.

| Factor | ‘lower noise’ group | ‘higher noise’ group |
| --- | --- | --- |
| External Noise | *** | *** |
| Improvements within-session | *** | * |
| Learning between the first day and the last day (start of the session) | n.s | ** |
| Learning between the first day and the last day (end of the session) | n.s | ** |
| Deteriorations between the end of the previous session and the beginning of the next session | *** | n.s |

